# Connectome quality converges predictably to reveal optimal stopping points during proofreading

**DOI:** 10.64898/2026.06.30.735414

**Authors:** Hannah Martinez, Jordan K. Matelsky, Daniel Xenes, Katharine Merfeld, Cara J. Cavanaugh, Patricia K. Rivlin, Cody J. Smith, Brock Wester

## Abstract

Volumetric electron microscopy (EM) has become a critical approach to generating high-resolution reconstructions of brain tissue. As the size of EM volumes increase, use of automated image segmentation within the reconstruction pipeline has become essential, although it generates errors that need correction. The proofreading and correcting of these errors has since become the dominant cost driver in the pipeline, but precisely estimating the sufficient number of proofreading edits to enable meaningful scientific analyses of the reconstructed neuronal networks remains a challenge. We present a fast, computationally inexpensive way to estimate the progress of a connectomic proofreading effort without requiring *a priori* knowledge of ground truth. We show that simple global graph invariants converge predictably to asymptotic limits with increasing numbers of proofreading edits, informing a quantitative “pencils down” criterion for proofreading completeness. We illustrate our method on two datasets in different stages of proofreading progress, a zebrafish spinal cord and the hemibrain *Drosophila melanogaster* dataset. Our method reduces the uncertainty associated with the planning and prioritization of proofreading activities and enables data owners to accurately predict and budget the amount of proofreading necessary for their scientific questions.

## Background

Advancements in volume electron microscopy (vEM) have been critical to reveal new architectural principles of tissues and organs ^1–3^. For example, analysis of vEM of brain tissue has created the field of nanoscale connectomics, which seeks to understand the synaptic connections of all neurons within a given sample of neural tissue ^4^. In recent years, nanoscale connectomes have grown dramatically in scale from single-neuron circuits to whole-brain volumes for the fly and cubic-millimeter scale volumes of the mouse and human brain ^5–9^. Modern EM reconstruction pipelines can segment teravoxel-to petavoxel-scale datasets automatically, but the resulting downstream data products, such as dense segmentations and connectome graphs, contain errors that can impact the accuracy of reconstructed circuits and downstream analysis^10, 11^. Before scientific analysis can be performed, dataset quality must be quantified, and merged cells, split fragments, and incorrect synaptic partners must be corrected through proofreading. For large datasets, the proofreading step dominates cost and time^11, 12^; for example, the recent MICrONS manual proofreading effort required 2.5 person years to proof-read about 5,500 undersegmented cells ^13^. Proofreaders required an average of 43 minutes, including training time, to identify, validate, and commit a single split edit ^13, 14^.

Despite this expense, there are few quantitative rules for deciding how much proofreading is sufficient to start subsequent network analyses^7, 14^. Numerous metrics exist for assessing reconstruction quality across progressive stages of EM dataset maturation, including those that measure imaging reliability, segmentation quality, and connectome accuracy ^15–17^. However, many of these metrics require access to painstakingly established manual ground truth on carefully curated subvolumes before measurement is possible. The gold standard is synaptic completeness, defined as the fraction of synapses in the dataset whose pre- and postsynaptic locations belong to fully proofread cells ^3, 18^. However, no large-scale EM dataset has achieved 100% synaptic completeness; two of the best proofread datasets in the field are the Flywire connectome at 44.7% and the hemibrain at 36%, and analysis showed that continued proofreading would strengthen existing pathways rather than reveal new ones^3, 7, 14^. Thus, the question remains – can allocation of proofreading resources be optimized with additional information on the proofreading trajectory?

Since graph theoretic analysis of the connectome is an important downstream goal of vEM, a possible approach to uncover a criterion for completeness for a dataset is to repeatedly analyze its graph theoretic properties, or graph invariants ^19^. Here, we propose a method to assess the proofreading state using global graph metrics and estimate the remaining edits necessary to reach a desired quality threshold. We discover that certain metrics, captured by simple and inexpensively computed invariants of the connectome graph, converge as proofreading progresses. By reanalyzing the graph at fixed intervals, we track how its invariants evolve and, by fitting monotone asymptotic models, forecast the remaining edits needed to meet a target network reconstruction quality. The prerequisites for this method are multiple connectomes computed at regular intervals over the proofreading history of a dataset, as well as the number of edits that were committed between connectome snapshots.

## Results

### Invariant convergence reveals a definition of done

We developed this approach using a serial block-face electron microscopy (SBEM) zebrafish spinal cord dataset^20^ that has received about 5,300 hours of manual merge-focused proofreading to improve errors in the image segmentation (**Fig. 1**). At the time of analysis, 2955 segmentation IDs have been identified as fully proofread cells. While proofreading is still ongoing, significant progress has been made towards achieving a level of synaptic completeness suitable for analysis (**Fig. 2**). We reasoned that by using the edit history, we could identify a proofreading checkpoint at which the properties of the global connectome graph would reduce their rates of change sufficiently enough that diminishing returns would no longer be worth the effort to continue proofreading. To measure this, we compute a set of 15 global graph invariants which measure qualities like shape, centrality, and clustering behavior on the synaptic connectome at eleven evenly spaced intervals in the proofreading changelog (**Fig. 3**). Generating eleven connectomes required significant computational resources, so we developed a new Python package, *Cloudome*, to distribute the process of connectome generation across multiple compute nodes (see **Methods** for more). In-variants that are computationally expensive to compute are approximated. We find that certain invariants tend to exhibit predictable convergence toward stable asymptotes (**Table 1**). This suggests that proofreading progress generates diminishing returns and indicates that forecasting the remaining effort left on a proofreading effort is possible using these invariants as proxies, even when the true error rate is unknown. After computing the number of edits at which each invariant will reach 50%, 80%, 90%, 95%, and 99% of its theoretical limit according to its respective fitted curve, we compute the mean and standard deviation of all converging invariant curves to define a composite metric for when the dataset as a whole will reach its theoretical limit (**Fig. 4**). For the zebrafish spinal cord, we find that six invariants will converge after approximately 71,000 edits, with an upper limit of 89,000 edits necessary to enable graph analysis on the global properties of the connectome. We can also estimate that the proofreading effort is at least 85% complete based on the 50,000 edits committed to the dataset so far. To remain conservative, we recommend that proofreading continue until the upper limit of 89,000 edits is reached, and that graph invariants continue to be recomputed at regular intervals. As more proofreading is performed and more metrics converge, the estimate may change.

**Figure 1.**
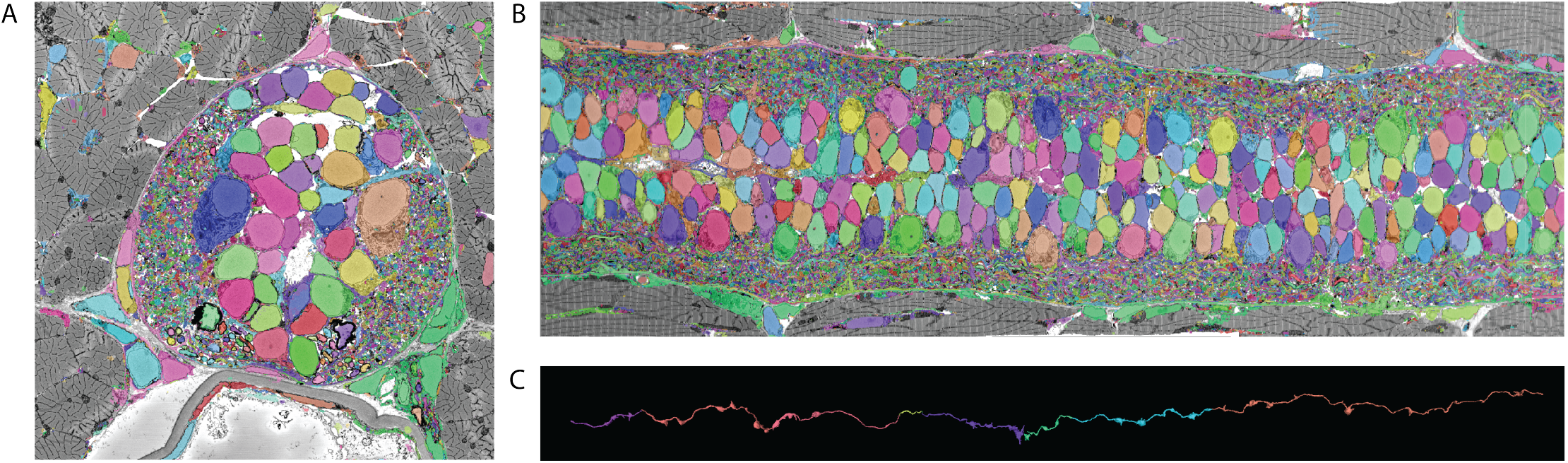
Zebrafish spinal cord electron microscopy and segmentation. **A-B**. The zebrafish spinal cord transverse view and lateral view. **C**. An axon that was progressively traced by a proofreader. Colors represent individual segments that were merged.

**Figure 2.**
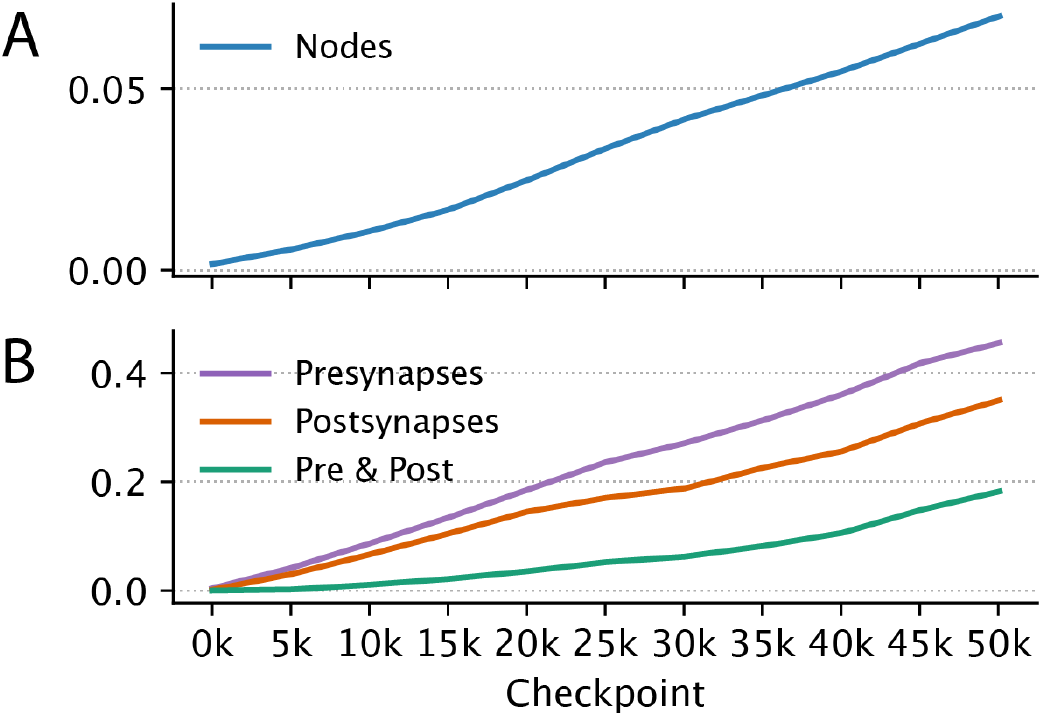
Synaptic completeness of the zebrafish spinal cord over time. **A)** Fraction of connectome nodes that correspond to cells that have been 100% proofread. **B)** Fraction of pre, post, and complete synapses belonging to nodes that have been 100% proofread. The synaptic completeness (green) approaches 20%.

**Figure 3.**
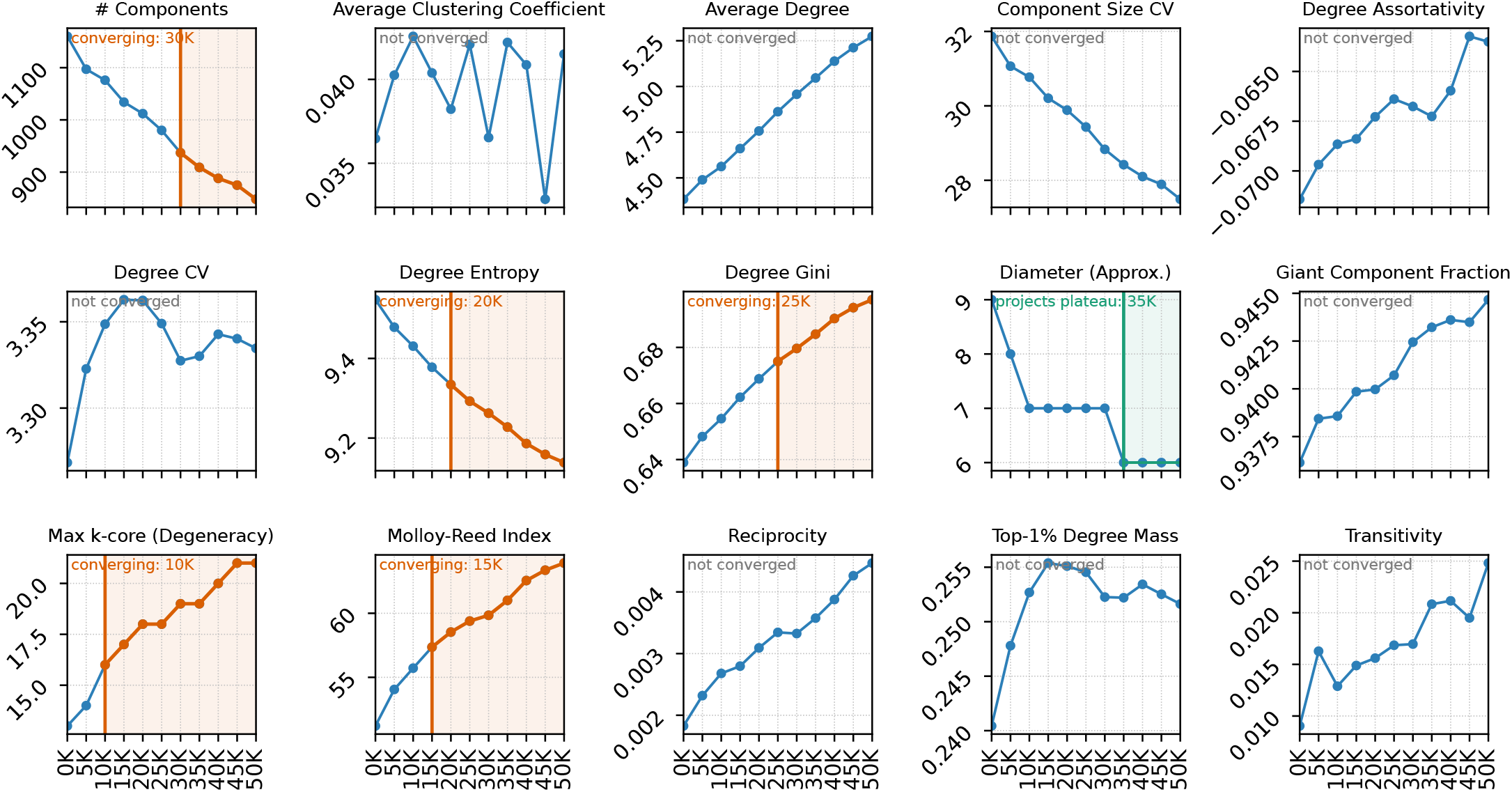
Connectome asymptotic invariant trajectories. The horizontal axis represents the number of edits on the zebrafish dataset. Each graph depicts the evolution of a different graph invariant computed on the connectome after the corresponding number of edits. An asymptotic curve is fit to each graph invariant plot. One invariant converges (green), while five more show converging behavior with forecastable limits (orange). Please refer to **Table 2** for a definition and discussion of each graph invariant.

**Table 1.** Zebrafish connectome invariant behavior. For each invariant showing converging behavior, we assume that its projected limit is its final value, then calculate the points at which it reaches *n*% of its final value.

| Invariant | Projection basis | Final value | 50% edits | 80% edits | 90% edits | 95% edits | 99% edits |
| --- | --- | --- | --- | --- | --- | --- | --- |
| diameter_approx | plateau signal | 6 | 7,500 | 32,000 | 33,500 | 34,250 | 34,850 |
| kcore_max | monotone tail | 22 | 17,500 | 41,000 | 50,760 | 55,760 | 67,370 |
| molloy_reed_index | monotone tail | 64.7838 | 17,707 | 38,424 | 45,562 | 52,407 | 68,555 |
| degree_entropy | monotone tail | 9.0844 | 22,201 | 41,676 | 52,401 | 63,230 | 88,373 |
| degree_gini | monotone tail | 0.704102 | 22,184 | 40,973 | 51,217 | 61,494 | 85,355 |
| num_components | monotone tail | 807.208 | 24,437 | 43,936 | 51,915 | 61,059 | 82,290 |
| Mean edits | — | — | 18,588 | 39,668 | 47,559 | 54,700 | 71,132 |
| Std edits | — | — | 5,554 | 3,788 | 6,680 | 9,865 | 18,080 |

**Figure 4.**
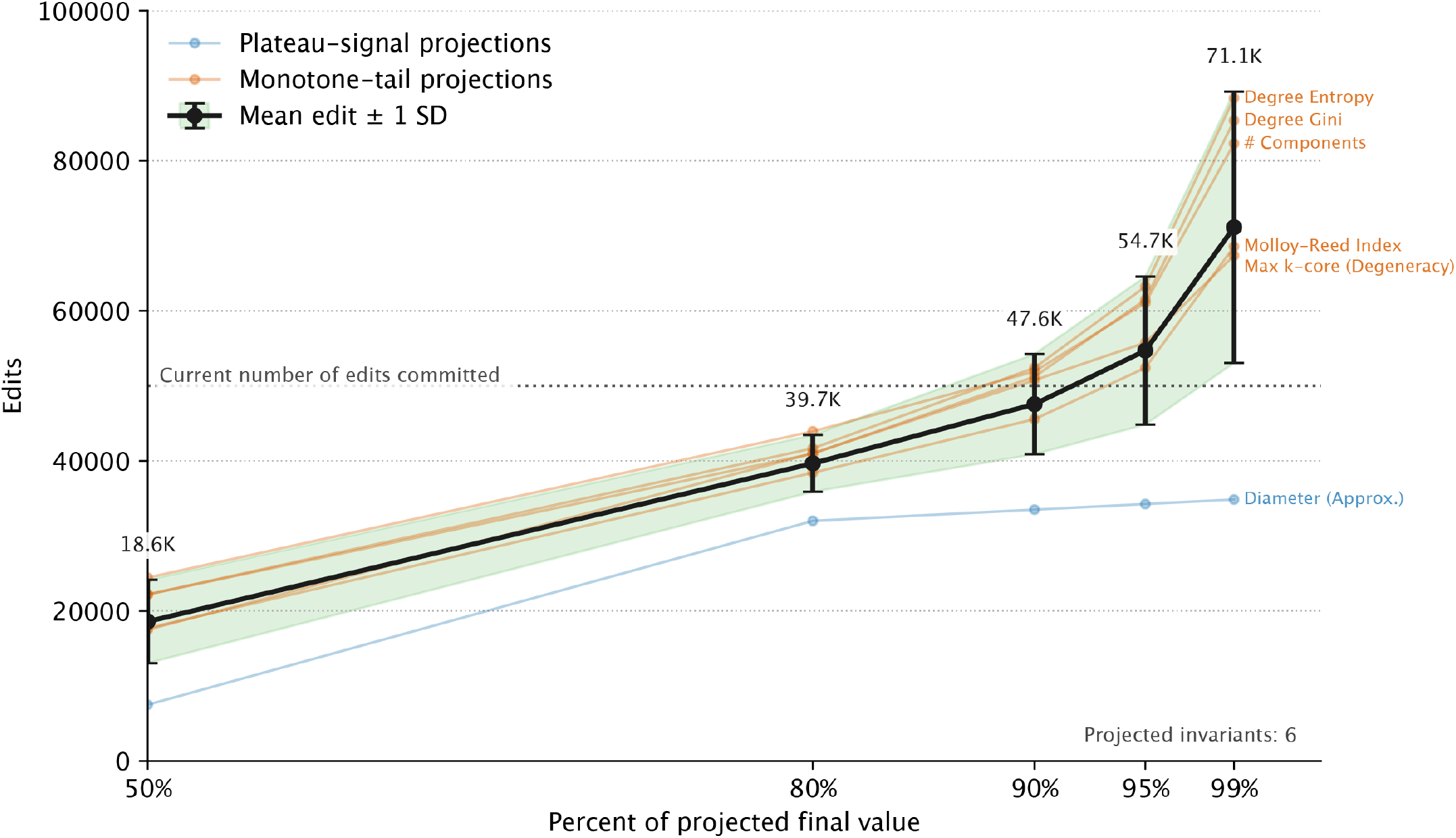
Combining Zebrafish graph invariant asymptotes to estimate the total proofreading progress. We use the progression towards invariant asymptotic behavior as a proxy for measuring the increase in dataset reliability as proofreading progresses. Since 50,000 edits have been committed to the dataset, we estimate that invariants have reached at least 85% of their projected values. This graph serves as a visual quantification of the diminishing returns that will be gained from continued proofreading.

### Validation on the hemibrain

In order to show that our method is generalizable across datasets, we replicated our asymptotic analysis on the hemibrain *Drosophila melanogaster* dataset, another community dataset whose image segmentation has undergone extensive proofreading.^3^. The hemibrain is a larger dataset with over 20,000 fully proofread neurons, and it used a more diverse set of proofreading strategies, including periods of focused merging, periods of focused splitting, and automated edit application. Thus, the hemibrain represents a larger and more complex dataset on which to validate our method. To do this, we pulled 11 connectivity snapshots spanning the available hemibrain proofreading history, representing a total of 1,732,048 edits, and replicated our analysis by measuring the asymptotic behavior of the same 15 graph invariants. We find that the trajectories of 12 graph invariants converge towards asymptotic limits, and that when these 12 are combined, the composite metric is consistent with over 95% completeness (**Fig. 5, 6**). While the asymptotic behavior of the hemibrain graph invariants is most obviously different from that of the zebrafish in scale, which can be attributed to the proofreading history being two orders of magnitude longer, we also observe that some invariants differ in trend direction. Since connectomics preprocessing methods vary widely across variables such as segmentation approach, supervoxel size, and agglomeration strategy, this is not surprising. We expect each dataset to exhibit a unique starting set of invariants that will trend upwards, downwards, or oscillate towards a true value as proofreading continues. We also observe that the composite metric has a larger standard deviation, reflecting that twice as many invariants show convergence and are included in the calculation. We believe this is so because some invariants converged earlier than others; thus, we continue to recommend that it be interpreted conservatively, with edit numbers on the higher end of the range reflecting the true number of edits required to achieve completeness.

**Figure 5.**
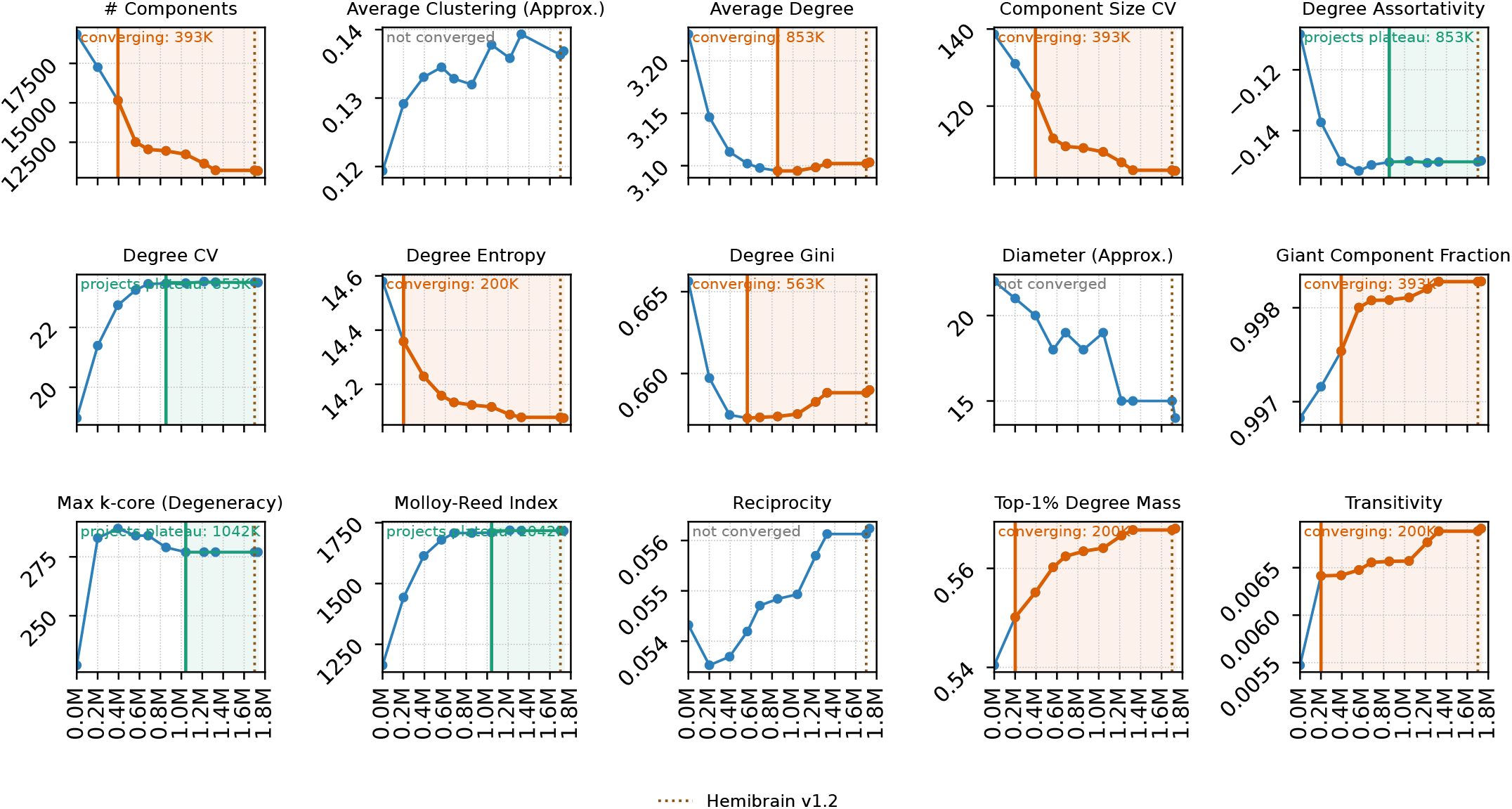
The hemibrain’s connectome asymptotic invariant trajectories. The horizontal axis represents the number of edits associated with each checkpoint. Four invariants converge (green), while eight more show converging behavior with forecastable limits (orange). The dotted line represents the true number of completed edits, 1,732,048. With twelve fitted curves, the hemibrain shows twice as much asymptotic behavior as the zebrafish. Please refer to **Table 2** for a definition and discussion of each graph invariant.

**Figure 6.**
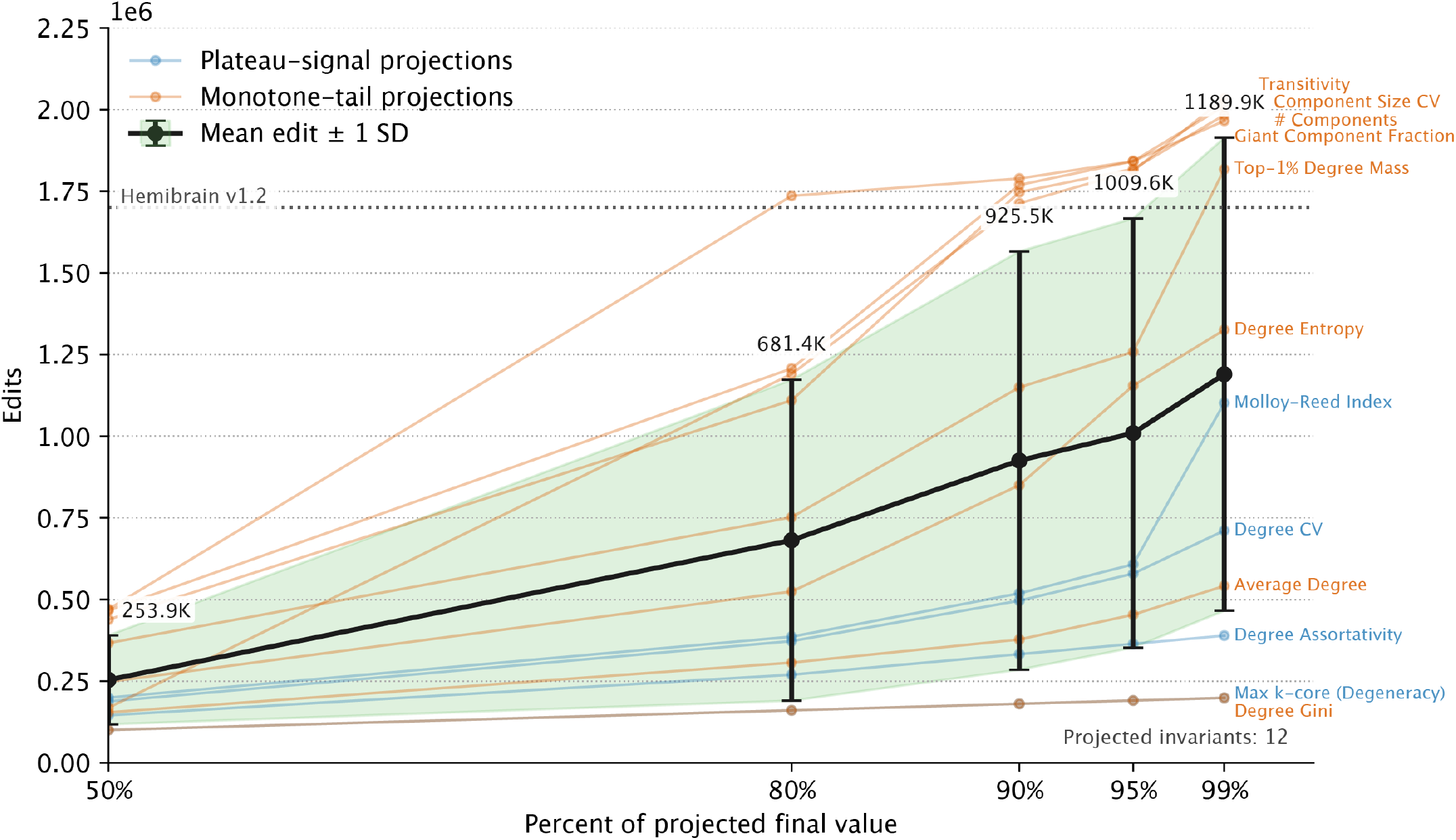
Combining the hemibrain’s graph invariants shows our composite metric on a dataset whose formal proofreading effort is finished. Since the hemibrain proofreading effort has been finalized, this graph demonstrates correspondence between our method’s assessment of completeness and a more qualitative consensus view. The dotted line shows the true number of completed edits, consistent with graph invariants having surpassed 95% of their projected values. Through this graph, we do not claim the hemibrain is “complete” beyond claims made in the original hemibrain paper; however, we do claim that continued proofreading would change the global network statistics very little.

### Forecasting remaining effort and stable values

We reasoned that by estimating asymptotes early and using the number of edits as a proxy for human labor (and thus cost), we could determine the necessary resources required to reach a desired completeness threshold. This level of precision in labor estimates would greatly improve the ability to schedule and budget the completion of connectomic reconstruction and subsequent scientific studies. Additionally, forecasting before proofreading is complete allows for estimations of the final values of each graph invariant (**Table 1**). We sought to understand the emergence and stability of these estimates by simulating the growing availability of checkpoint information over time. To do so, we iteratively removed each latest proofreading checkpoint and recalculated our milestones to determine at what point enough proofreading had occurred to accurately produce a forecast for remaining effort (**Fig. 7**). We found that for the zebrafish, the asymptotic fits of the component graph invariants became stable around the 40,000 edit checkpoint; for the hemibrain, the fits became stable around 1.2 million edits. If we consider our estimate of 71,000 edits on the zebrafish to represent 99% proofread, as per our previous analysis, then we have found that the zebrafish stopping point became forecast-able when around 56% of the total necessary proofreading was complete. For the hemibrain, 1.2 million edits represents about 70% of the total reported proofreading history, and we note that some proofreading took place before edit history was tracked in DVID, so this ratio may be an underestimate^21^. We note that the hemibrain proofreading effort utilized more diverse proofreading strategies, such as periods of focused splitting and automated methods, so a slower convergence pattern is not surprising. Because these two milestones are so different, we emphasize that more work will be needed to determine how early in the proofreading process the necessary effort can be forecasted. For future proofreading, we recommend that **Fig. 7** graphs are generated early and often for each volume or subvolume to ascertain when estimates stabilize.

**Figure 7.**
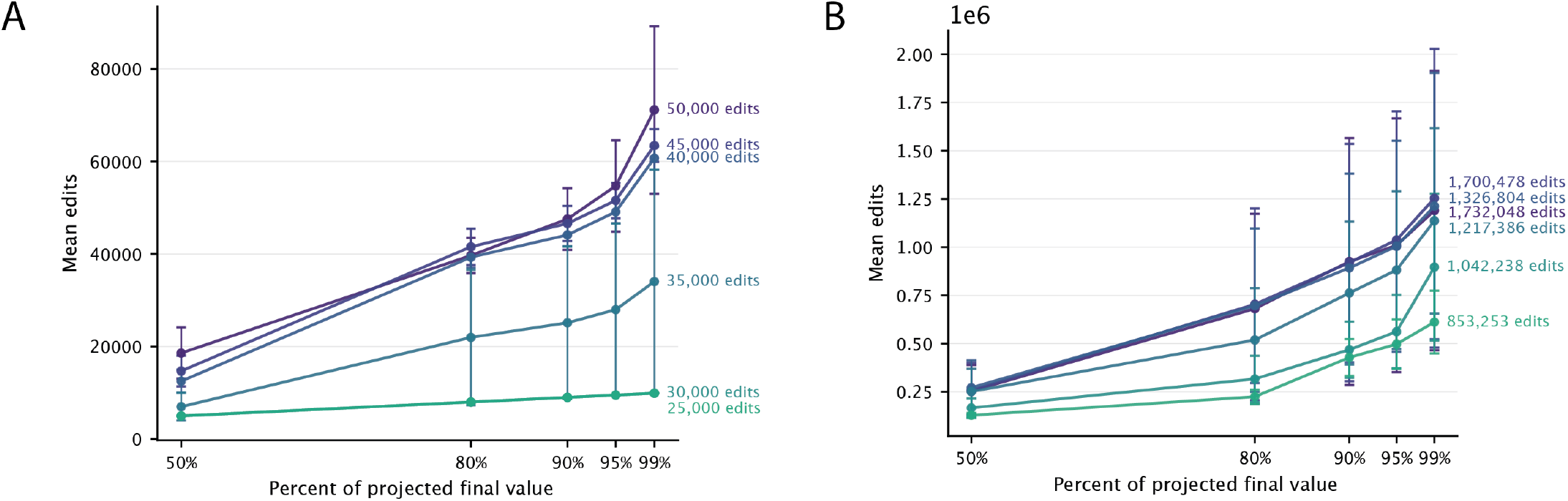
Estimating asymptotes early reveals the amount of effort remaining. Here we iteratively remove checkpoints to determine how early in the proofreading process an accurate forecast can be made. We find that for the zebrafish spinal cord **(A)**, the composite metric becomes stable around the 40,000 edit point, and for the hemibrain **(B)**, the invariant composite metric becomes stable around the 1.2 million edit point. We recommend generating these graphs early and often during proofreading, as the estimate will become more accurate as edits are committed.

Future work will address scaling up to larger, whole-brain volumes. Additional variables, such as number of synapses, supervoxel size, and anatomical region, may also affect the threshold at which the composite metric enables forecasting.

## Discussion

We have shown that the global graph invariants of an in-progress synaptic connectome have a convergent limiting behavior that provides a clear, quantitative stopping criterion for proofreading. Instead of relying on intuition or budget ceilings, teams can specify a tolerated error level and compute the remaining effort. We focus on global graph invariants because they do not require *a priori* knowledge of ground truth to calculate and because they directly measure properties of the synaptic connectome. By contrast, voxel-based metrics that quantify segmentation accuracy are standardized, but correlate imperfectly with the underlying network graph, while path-based metrics like expected run length (ERL) are informative but require painstakingly collected ground truth ^17, 22^. One limitation with the use of global graph invariants is that they function as proxy metrics in this analysis and cannot be relied upon to measure progress towards a specific scientific question beyond global graph analysis. We propose that future proofreading efforts use this method to determine a practical definition of “done” at which time a dataset can be released for community use, with additional targeted proofreading performed on a case-by-case basis depending on each scientific question that is answered in accompanying papers.

Recent work has increasingly highlighted the need for new methods that can both measure reconstruction quality and include a confidence interval on that measurement ^23^. We argue that tracking the evolution of graph invariants over successive proofreading states is attractive not because it replaces direct error measurement, but because it offers a method to detect stabilization in the connectome structure and therefore acts a proxy for dataset reliability at the global level. Importantly, our method should not be interpreted as measuring changes in the biological properties of the underlying circuit, which were fixed at the time the tissue was imaged. Rather, the observed trajectories reflect the human process of moving the measured graph toward the true values of those biological properties. We intend for our composite metric to work alongside other established metrics to produce a holistic view of the quality of a dataset, and expect that it can be used not just with manual workflows, but also with the growing collection of algorithms that automatically proofread EM volumes such as NEURD, PATHFINDER, and RoboEM to determine how many computational resources to allocate towards these tools^11, 12, 16^. Since we demonstrate the approach on two cropped volumes for which two different proofreading strategies were used, we also expect that it can be used flexibly within a single dataset to compare subvolume proofreading progress or to compare the effects of differing merging or splitting approaches utilized during different time intervals.

In summary, our proposed composite metric relies on simple graph invariants, making it fast (i.e., does not depend on explicit human evaluation), scalable (i.e., can be calculated with modest compute resources), repeatable (i.e., regularly run as proofreading progresses), and compatible with any reconstruction pipeline that captures an edit history or for which snapshots may be taken. We expect that this metric can apply to any vEM proofreading effort, including those still in progress and those that focus on subsets of segmentation or connectomic data.

## Acknowledgements

Research reported in this publication was supported by the National Institute Of Mental Health of the National Institutes of Health under Award Number R24MH114785 and by National Institute Of Neurological Disorders And Stroke of the National Institutes of Health under Award Numbers U01NS137250 and U24NS139927. The content is solely the responsibility of the authors and does not necessarily represent the official views of the National Institutes of Health. We also thank the University of Notre Dame and the Gallagher family for supporting this work. We thank Ariadne for providing resources and infrastructure for proofreading. We thank Stuart Berg at Janelia Research Campus for providing the checkpointed hemibrain connectomes necessary to validate our results. Finally, we thank Zach Wrobel, Katharine Merfeld (author), Andrea Colon Del Rio, Turner Long, Antonio Dolman, Graham Mercurio, Harry Durgin, Joseph Lee, Lukas Abraham, Sophia Tran, Ache Ekokobe, Bryce Miller, David Ke, Ethan Tam, Jacob Suresh, Katie Ford, Liam Sweeney, Brady Shin, Cara Cavanaugh (author), David Diaz, Chang Chau, Zachary Koh, and Ricky Avalos for their time spent proofreading the zebrafish spinal cord.

## Code and Data Availability

All code produced for this study will be made available upon publication.

## Methods

The zebrafish spinal cord dataset includes SBEM images, a supervoxelized cell reconstruction, synapse detections that distinguish presynaptic and postsynaptic regions, and a proofreading changelog^20^. Changelog entries consist of the two affected supervoxel IDs and whether they underwent a merge or split. In this dataset, the changelog is predominantly composed of merges, where two supervoxels that are part of the same cell are joined, with a small fraction of splits reversing erroneous merges. Manual-only proofreading has primarily taken place on Z slice 4974 in the center of the dataset. Proofreaders are assigned segmentation IDs that bisect this Z slice and are instructed to trace these IDs in both Z directions, performing merges between their designated ID and additional IDs that belong to the same cell. This region-focused proofreading approach means that some regions of the dataset are better proofread than others.

### Checkpointed reconstruction generation

Since multiple synapse graphs were not immediately available to us given the input data, we iteratively applied merge and split edits from the proofreading changelog to the machine-generated supervoxels, saving snapshots of the segmentation at fixed intervals. While synapse detections remained fixed throughout proofreading, the segmentation IDs underlying the synapse detections changed, causing the synapse graph to change. At every snapshot, we recomputed the synapse graph, yielding the evolution of the network structure as proofreading progressed.

### Workload distribution and parallelization

Since the scale of this volume requires a computationally scalable method that can be run successively and automatically, we developed a software package, *Cloudome*, which divides work into small tasks, distributes them on a high-performance computing cluster or a serverless cloud backend, and combines the results (**Fig. 8A**). *Cloudome* supports rapid generation of connectomes directly from segmentation and contains a flexible framework for chunkwise calculations useful for computing contactomes, cell volumes, and more.

**Figure 8.**
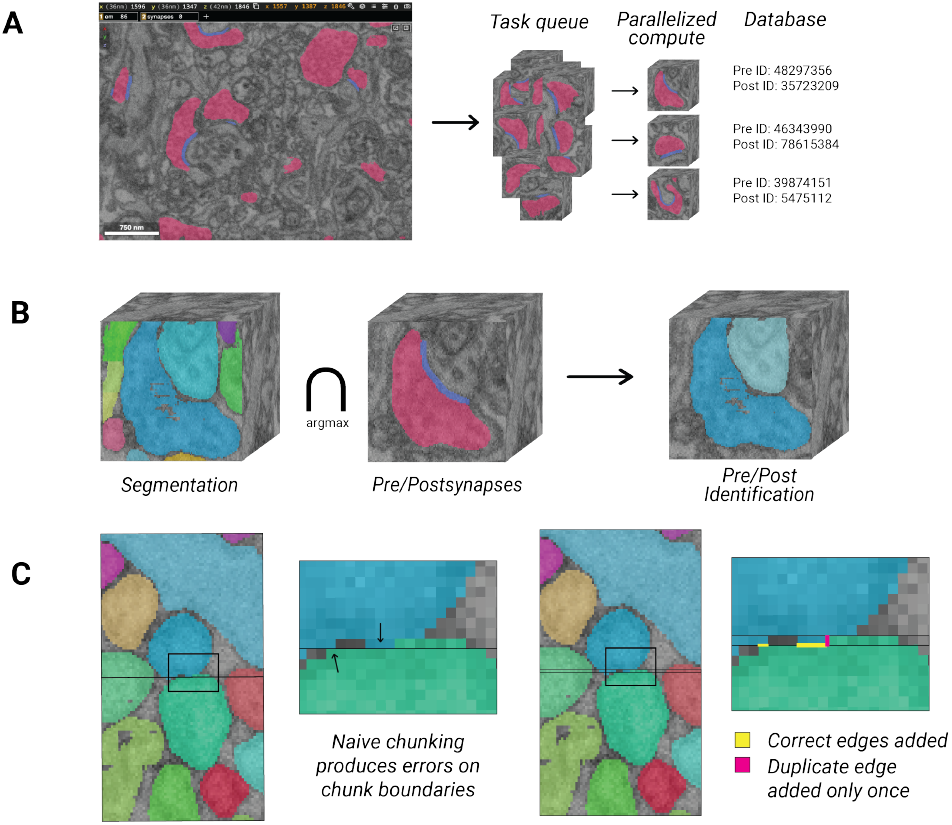
*Cloudome* makes task division and queueing easy. A) For a connectome task, an EM image in Neuroglancer helps the user identify individual synapses, with pink representing the presynapse and blue representing the postsynaptic density^26^. To generate a full connectome, *Cloudome* creates one pre- and postsynaptic partner identification task per synapse and schedules them on parallelizable architecture, then saves results in a database. B) One connectome task identifies the cell IDs participating in one synapse. Two registered cuboids of segmentation and synapse paint are compared to determine synaptic partners. These two data layers are prerequisites for a *Cloudome*-computed connectome. C) A contactome task is one kind of volumetric task that *Cloudome* supports. To compute one contactome task, the contacting surface area of adjacent segmentation IDs is computed for one cuboid of segmentation. A one voxel overlap on three out of six chunk faces is added to account for edge effects (yellow). Duplicated edges in the overlap regions (pink) are accounted for only once.

*Cloudome* splits connectome generation into one task per synapse, scheduled on the chosen parallel architecture. Each task requires a single synapse detection and a snapshot of segmentation overlaying the detection. The code then identifies the two cells that participate in that synapse by superimposing the synapse paint on the segmentation snapshot and selecting the segmentation IDs with maximal overlap (**Fig. 8B**). The task outputs are concatenated to produce the full history of the synapse graph.

For chunked tasks like cell contact and volumetric calculations, *Cloudome* subdivides 3D images from a data volume in precomputed format into chunks sized to the target machine. *Cloudome* supports both local datasets and datasets stored in cloud-based resources such as Google Cloud Platform and BossDB^24^. For serverless cloud infrastructure, we use 64*×*64*×*32 voxels; for a high-performance computing cluster, we use 1024*×*1024*×*1024 voxels. Chunks are processed independently to produce 3D measurements using connected-components-3d.^25^ These tasks scale linearly with volume.

### Edge effect compensation

“Chunkwise” processing as we describe here is subject to edge effect errors — cases where the properties of voxels on the boundary of a chunk are not properly computed because of the missing data outside the margin of the chunk context. Mitigation is most easily performed by computing an enlarged region with adequate margin to enable later cropping without compromising data integrity. Here we include a one-voxel overlap to ensure all edges are captured and subtract duplicated measurements appropriately (**Fig. 8C**).

### Invariant computation and model fitting

At each proofreading checkpoint, we compute 15 graph invariants on the full synaptic connectome (**Fig. 3, 5**). Invariants that are computationally expensive to compute are approximated. Each invariant is evaluated on the directed or undirected graph *G* = (*V, E*), where *V* denotes neurons and *E* denotes synaptic edges. For calculations involving connected components, we convert the directed graph *G* to an undirected graph before computation. In **Table 2**, we briefly define each metric formally and discuss how its value might change during proofreading.

**Table 2.** Global graph invariants used to track proofreading progress. These invariants summarize global properties of the reconstructed graph and are expected to converge as proofreading progresses.

| Invariant | Definition | Description | Expected change during proofreading |
| --- | --- | --- | --- |
| Average clustering coefficient | $\frac{1}{\vert V\vert } \sum_i C_i$ where for node $i$, $C_i = \frac{2T_i}{k_i(k_i-1)}$, and $T_i$ is the number of triangles incident to $i$ and $k_i$ is degree | Measures the tendency of a node's neighbors to connect with one another. | Converges as spurious local triangles are removed and local connectivity patterns stabilize. Can be approximated on a subset of nodes. |
| Transitivity | $\frac{3 \times \text{number of triangles}}{\text{number of connected triples}}$ | Measures the global fraction of closed triads in the graph. | Stabilizes as topological errors are corrected. |
| Diameter | $\max_{i,j \in V} d(i,j)$<br>where $d(i,j)$ is shortest-path distance in the largest connected component | Quantifies the longest minimal path through the network. | Expected to increase as erroneous merges are resolved. For large graphs, this metric can be approximated by running on a subset of $V$. |
| Molloy–Reed index | $\kappa = \frac{\langle k^2 \rangle}{\langle k \rangle}$, where $k$ denotes degree and $\langle \rangle$ indicates averaging over nodes | Governs the existence and robustness of a giant connected component in random graph theory. | Stabilizes as merged hubs and fragmented nodes are corrected. |
| Degree assortativity | $r = \rho(k_i, k_j)$, or the Pearson correlation coefficient for degrees of nodes at either end of an edge | Measures whether high-degree nodes preferentially connect to other high-degree nodes. | Stabilizes as proofreading corrects distortions in hub structure. |
| Degree entropy | $H = -\sum_k p_k \log p_k$, where $p_k$ is the fraction of nodes with degree $k$ | Quantifies heterogeneity in the degree distribution. | Stabilizes as artificial degree inflation and fragmentation are reduced. |
| Max $k$-core / degeneracy | Largest $k$ for which the graph contains an induced subgraph where every vertex has degree at least $k$<sup>27</sup> | Quantifies the density of the graph. | Stabilizes as the dense core structure of the reconstructed graph approaches its final value. |
| Degree Gini | $\frac{2 \sum_{i=1}^n i y_i}{n \sum_{i=1}^n y_i} - \frac{n+1}{n}$ | Measures inequality in node degree. A Gini coefficient of 0 means all nodes have the same degree, while a high value means a small number of nodes have very high degree. | Converges as artificial hubs and fragmented nodes are corrected. |
| Number of components | Number of connected components in the graph | Counts isolated subgraphs and unresolved fragmentation. | Expected to decrease as fragments are connected into the main graph. |
| Average degree | $\frac{2\vert E\vert }{\vert V\vert }$ | Measures the average number of connections per node. | Typically decreases during split-heavy periods and increases during merge-heavy periods. |
| Component size coefficient of variation | $CV = \frac{\sigma}{\mu}$, computed over the component size distributio. | Measures variability in connected-component sizes. | Stabilizes as fragmented components merge into larger structures. |
| Degree coefficient of variation | $CV = \frac{\sigma}{\mu}$, computed over the degree distribution | Measures variability in node degree. | Stabilizes as artificial degree inflation and deflation are corrected. |
| Giant component fraction | Fraction of vertices belonging to the largest connected component. | Measures how much of the graph belongs to the dominant connected component. | Expected to increase as isolated fragments join the main graph. |
| Reciprocity | $\frac{\text{number of reciprocal edges}}{\vert E\vert }$ | Measures the fraction of directed edges that are bidirectional. | Stabilizes as erroneous synaptic partners and segmentation artifacts are corrected. |
| Top 1% degree mass | $\frac{\text{degree}(V')}{\text{degree}(V)}$, where $V'$ is the 1% of vertices with highest degree | Measures how concentrated edges are among the highest-degree nodes. | Stabilizes as artificial hubs are corrected and true hub structure is recovered. |

### Model fitting and asymptote estimation

Each invariant is modeled as a function of cumulative edit count (**Table 1**). We fit families of strictly monotone asymptotic models (inverse power laws, exponential decay, and low-order polynomials). Model selection uses Bayesian Information Criterion (BIC),

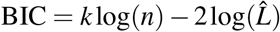

where *k* is the number of parameters, *n* the number of checkpoints, and 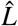 the maximized likelihood. Simpler convergent models are preferred when supported by the data. The fitted asymptote provides both an estimate of the final invariant value and a forecast of the remaining proofreading required to reach a specified fraction of convergence.

